# In silico analysis of long non-coding RNAs in medulloblastoma and its subgroups

**DOI:** 10.1101/783092

**Authors:** Piyush Joshi, Ranjan J. Perera

## Abstract

Medulloblastoma is the most common malignant pediatric brain tumor with high fatality rate. Recent large-scale studies utilizing genome-wide technologies have sub-grouped medulloblastomas into four major subgroups: wingless (WNT), sonic hedgehog (SHH), group 3, and group 4. However, there has yet to be a global analysis of long non-coding RNAs, a crucial part of the regulatory transcriptome, in medulloblastoma. Here, we performed bioinformatic analysis of RNA-seq data from 175 medulloblastoma patients. Differential lncRNA expression sub-grouped medulloblastomas into the four main molecular subgroups. Some of these lncRNAs were subgroup-specific, with a random forest-based machine-learning algorithm identifying an 11-lncRNA diagnostic signature. We also validated the diagnostic signature in patient derived xenograft (PDX) models. We further identified a 17-lncRNA prognostic model using LASSO based penalized Cox’ PH model (Score HR= 13.6301, 95% CI= 8.857-20.98, logrank p-value=< 2e-16). Our analysis represents the first global lncRNA analysis in medulloblastoma. Our results identify putative candidate lncRNAs that could be evaluated for their functional role in medulloblastoma genesis and progression or as diagnostic and prognostic biomarkers.

## Introduction

Medulloblastoma (MB), characterized as WHO group IV, represents the most common malignant pediatric central nervous system (CNS) tumor (Diamandis and Aldape, 2018; Louis et al., 2007; Northcott et al., 2019; Ostrom et al., 2017), representing 9.2% of all pediatric brain tumor cases (Millard and De Braganca, 2016; Ostrom et al., 2017) and roughly 500 new cases each year in the US. MBs originate in the cerebellum and share molecular signatures with embryonic cerebellar lineages, with metastasis sites commonly include parts of the brain, spinal cord, and, rarely, to extraneural sites (Dufour et al., 2012; Kondoff et al., 2015; Vladoiu et al., 2019).

Commonly used treatment strategies for MB include maximal safe surgical resection, radiotherapy, and chemotherapy, which are poorly tolerated by pediatric patients who are usually under seven years of age (Smee et al., 2012). Appropriate treatment selection depends upon the clinical subgroup, stage, extent of resection, location, and the patient's ability to withstand treatment (De Braganca and Packer, 2013). In efforts to improve therapeutic outcomes, combined genetic and epigenetic approaches have refined MB classification into four clinically and molecularly distinct subgroups: wingless (WNT), sonic hedgehog (SHH), group 3, and group 4 (Cavalli et al., 2017; Northcott et al., 2017; Northcott et al., 2012). The molecularly distinct group also present distinct clinical, pathological and prognostic feature; for example, WNT and group 3 patients represent prognostically best and worst subgroups, respectively, among MBs (for a detailed review on MB pathology, we recommend following comprehensive reviews: Millard and De Braganca, 2016; Northcott et al., 2019). Despite these significant diagnostics advances, MB remains deadly for many patients, with a ~30% fatality rate. Further, even successful eradication of the tumor often results in a deteriorated overall quality of life due to side effects including organ dysfunction, neurocognitive impairment, endocrine disabilities, and secondary tumors (De Braganca and Packer, 2013; Martin et al., 2014; Palmer et al., 2007; Wang et al., 2018a). In addition, even with advances in molecular classification, group 3 and group 4 tumors represent heterogeneous groups that continue to make management challenging. There is an urgent need to identify the underlying molecular mechanisms in these subgroups to drive precision medicine-based approaches, improve quality of life, and increase our understanding of MB in general.

Long non-coding RNAs (lncRNAs) represent a major part of the transcribed genome that do not code for functional proteins. LncRNAs are more than 200 nucleotides in length and are transcribed by RNA polymerase II. While previously labelled as transcriptional “noise”, it is now understood that lncRNAs are functional and play important roles in cellular physiology, development, and disease progression. In humans, there are at least three times as many lncRNAs as protein-coding genes (Iyer et al., 2015). Although the precise roles of the vast majority of identified or predicted lncRNAs remain unknown (Iyer et al., 2015), they are increasingly recognized as being involved in *cis* or *trans* interactions regulating gene expression in the nucleus and protein interactions in the nucleus and cytoplasm. Some of the functionally diverse roles of lncRNAs include transcriptional silencing (e.g., *XIST* (Sahakyan et al., 2018)), enhancers by regulating three-dimensional chromosomal structure to strengthen interactions between enhancers and promoters (e.g., *LUNAR1* (Trimarchi et al., 2014)), and as microRNA sponges that sequester microRNAs from their target sites (e.g., *SNHG7* (Shan et al., 2018)). LncRNAs can also act as scaffolds for protein-protein and protein-nucleic acid interactions (Long et al., 2017). They are potential biomarkers and therapeutic targets in cancer, with several lncRNAs now studied for their oncogenic or tumor suppressor potential in several cancer types through their regulation of the cell cycle, cell death, senescence, metastasis, immunity, and cancer cell metabolism (Huarte, 2015).

LncRNAs are also known to be implicated in CNS tumors including various type of gliomas (Reon et al., 2016; Zhang et al., 2012; reviewed in Pop et al., 2018). However, there has yet to be a genome-wide study of MB to identify dysregulated lncRNAs. With this aim, we analyzed the transcriptomic profiles of 175 MB patients to map lncRNA expression profiles and identify subgroup-specific lncRNAs. We show that the MB lncRNAome exhibits significant heterogeneity that corresponds to the molecular subtypes. Using a random forest-based machine-learning algorithm, we identify lncRNA signatures that could improve on present diagnostic approaches, while penalized Cox-PH regression identifies prognostic lncRNAs. Taken together, our analysis identifies candidate lncRNAs with subgroup-specific activity in MB and with diagnostic and prognostic value.

## Materials and methods

### Datasets

Raw FASTQ files for RNA-seq data corresponding to 175 MB patients (referred to as the ICGC dataset, accession number EGAD00001003279) belonging to four subgroups were downloaded from the European Genome-Phenome Archive (EGA, http://www.ebi.ac.uk/ega/) after obtaining approval from the Institutional Review Board (IRB) (**Table S1**) (Northcott et al., 2017). The RNA-seq data were generated in four different studies in International Cancer Genome Consortium (ICGC) PedBrain tumor project (Jones et al., 2012; Lin et al., 2016; Northcott et al., 2017; Northcott et al., 2014). Pre-analyzed microarray expression datasets from 763 patients belonging to the four medulloblastoma subgroups were obtained from the study published by Cavalli et al. (referred to as the MAGIC dataset) (Cavalli et al., 2017).

### RNA-seq library preparation

RNA sequencing for patient derived xenograft (PDX) samples was undertaken at the Genetic Resources Core Facility at the Johns Hopkins University, School of Medicine, Baltimore, MD. Before sequencing, total RNA was extracted from PDX cell pellets using the Direct-zol RNA miniprep kit (R2060, Zymo Research, Irvine, CA), with subsequent quantification using Nanodrop (Thermo Fisher Scientific, Waltham, MA) and quality assessment with the Agilent Bioanalyzer Nano Assay (Agilent Technologies, Santa Clara, CA). Strand specific RNA-seq libraries were constructed using the Illumina TruSeq Stranded Total RNA Library preparation Gold kit (20020598, Illumina Inc., San Diego, CA) as per the instructions. The quality and quantity of the libraries were analyzed using the Agilent Bioanalyzer and Kapa Biosystems qPCR (Sigma Aldrich, St. Louis, MO). Multiplexed libraries were pooled, and paired-end 50 base-pair sequencing was performed on a NovaSeq6000. PDX RNA-seq data is available at the Gene Expression Omnibus (GEO) Accession Number GSE134248.

Libraries for ICGC RNA-seq data in original publications were prepared using a strand specific method (Truseq kit or modified method), sequenced on Illumina HiSeq2000 platform for 2X51 cycles.

### Quantitative RT-qPCR analysis

Expression of 9 candidate diagnostic lncRNAs were validated in PDX samples. RNA was isolated from PDX samples as described above. To design the primers (Table S2), qPCR primer pairs covering exon-exon boundaries (in multi-exonic transcripts) for most abundant gene transcript (as analyzed from RNA-seq data of PDX samples) or most common exonic boundary (in case more than one transcript showed comparable expression) were designed. More than one pair of primers were designed with the pair showing an exponential curve in qPCR (for each candidate) used for further analysis. Expression of *ACTB* gene was used to obtain normalized Ct values as only *ACTB* showed least variant expression among housekeeping genes in MB patients both in RNA-seq and microarray dataset.

### RNA-seq alignment, quantification, and differential gene expression analysis

Raw FASTQ files (from both public data and in house generated RNA-seq of PDX samples) were quality checked for adapter contamination using FASTQC (v0.11.9). FASTQ files containing adapter sequences were trimmed by running through Trim Galore (v0.5.0) in default mode. The reads were mapped to the GRCh38/hg38 human genome assembly p12 (release 28, www.gencodegenes.org) using HISAT2 (v2.1.0) and annotated using the corresponding release version GENCODE comprehensive annotation file and LNCipedia 5.2 high confidence set annotation file. LNCipedia high confidence set lncRNAs, with Ensemble ids overlapping with GENCODE annotated set, were removed from GENCODE comprehensive annotations and kept in LNCipedia annotation file to avoid duplication of lncRNAs. Mapped reads were quantified using StringTie (v1.3.6) to obtain TPM values, which were converted to read counts using the prepDE.py script (provided in online StringTie manual webpage). For variance-stabilized normalized reads and differential gene expression analysis, reads counts were processed with *DESeq2 (v1.24.0)* in R 3.6.1 (Love et al., 2014).

### Consensus clustering

Variance-stabilized expression levels of the top 10,000 variant lncRNAs determined from standard deviations of read counts normalized to library size were used as input to perform 1000 permutations of k-means-based consensus clustering using *ConsensusClusterPlus (v1.48.0)* R package (Wilkerson and Hayes, 2010).

### Co-expression module detection and trait correlation analysis

Variance-stabilized expression of top 5000 variant lncRNAs was used to obtain a weighted correlation network using the *WGCNA (v1.68)* R package (Langfelder and Horvath, 2008). The correlated lncRNA cohorts were associated with MB subgroups using the module-trait correlation algorithm as described (Langfelder and Horvath, 2008).

### Random forest model

Subgroup-specific diagnostic models were obtained by performing variable selection using expression of differentially expressed lncRNAs/ protein coding genes (PCGs) and the *randomForest (v4.6-14)* R package (as described in (Mehrian-Shai et al., 2007)). For all models, variance-stabilized expression of differentially expressed lncRNA/PCG genes were used as variables to obtained models to classify patients into known subgroups. For the lncRNA model distinguishing SHH, group 3, and group 4, patient samples were divided into a 60% training set and 40% tuning set. Only differentially expressed (|logFC| >1.5, padj <0.01) lncRNAs genes between SHH, group 3, and group 4 were used to classify patients into known subgroups. The training model was used to find important genes ranked based on the “mean decrease accuracy” parameter. Low ranking genes with high expression correlation (>0.80) to high ranked genes were discarded. Gene combinations based on the final ranked list were used in the tuning model to find the minimum number of genes resulting in the minimum or comparatively lower error rate in the tuning set. A similar approach was used to distinguish WNT from the other subgroups. For the training set, all WNT samples were combined with 60% samples from SHH, group 3, and group 4 samples. For the tuning set, all WNT samples were combined with 40% training set the remaining subgroups. To distinguish group 3 and group 4, a similar 60%-40% training-tuning model was adapted for classification.

A random forest-based model for PCGs distinguishing WNT, SHH, group 3, and group 4 was obtained as described above for lncRNAs. To validate the protein coding model, expression levels of the obtained signature genes were used to classify patient samples from the MAGIC dataset using the random forest model and tSNE plots.

Receiver operating characteristics (ROC) curves and area under the ROC curve (AUC) values for one versus rest comparisons were computed based on a generalized linear model-based fit of subgroup identity with normalized gene expression levels of signature genes as the variable using the *pROC (v1.61.1)* R package.

### Transcriptional network inference

A transcriptional inference network for putative regulation between candidate lncRNAs and transcription factors was obtained using *minet (v3.42.0)* R package (Meyer et al., 2008). Regulatory interaction measures were obtained based on the network obtained from CLR-, arcane-, and mrnet-based models. Only edge connections predicted in all three approaches were analyzed further for first neighbor connections of transcription factors with each candidate lncRNA.

### Survival analysis

Expression of 621 lncRNAs (identified based on Ensembl biotype annotation) included in the MAGIC microarray dataset for 612 patients with survival information belonging to the four subgroups was used as input to find prognostic signatures using penalized Cox’s proportional hazard (Cox-PH) model with the LASSO (α =1) penalty using *glmnet (v2.0-18)* R package (Tibshirani, 1997). Smallest mean squared error and *lambda.min* was selected from 100 random runs of the model fitting with 10 fold cross validation in each run. Variables (lncRNAs) with non-zero coefficients associated with the *lambda.min* were selected as the prognostic model. Stability of the obtained model was verified by performing 1000 bootstraps on the data using *BootValidation (v0.1.65)* R package. A risk score was calculated for each patient by summing the value of the product of normalized expression and penalized Cox-PH coefficient of a candidate lncRNA with that of all other candidate lncRNAs. Kaplan-Meier analysis was conducted using the obtained risk score of a 17-lncRNA prognostic model for the MAGIC dataset with survival information and 74 patients belonging to four subgroups with survival information in the ICGC RNA-seq dataset.

## Results

### Long non-coding RNA signatures of medulloblastoma and subgroup-specific lncRNA enrichment

To determine genome-wide expression profiles of lncRNAs in MB, we analyzed RNA-seq data from 175 patient samples obtained from the ICGC PedBrain tumor dataset. For a comprehensive lncRNA annotation, we chose a combination of the GENCODE (Frankish et al., 2019) and LNCipedia (Volders et al., 2019) datasets: GENCODE represents the largest manually curated lncRNA dataset and LNCipedia contains the maximum number of high fidelity predicted lncRNA genes (Uszczynska-Ratajczak et al., 2018). The expression of 52,128 unique lncRNAs and 19,033 protein-coding genes (PCGs) were quantified (**Fig 1A**) in 18 WNT MBs, 46 SHH MBs, 45 group 3 MBs, and 66 group 4 MBs. To better understand the role of lncRNAs in different MB clinical and molecular subgroups, we investigated correlations between lncRNA expression and subgroup type. First, we performed consensus clustering using the top 10,000 highly variant lncRNAs, which clustered the MBs into four different groups that highly overlapped with the known molecular subgroups (**Fig 1B, Fig S1, Table S3**), suggesting that lncRNAs could contribute to subgroup-specific traits.

**Figure 1.**
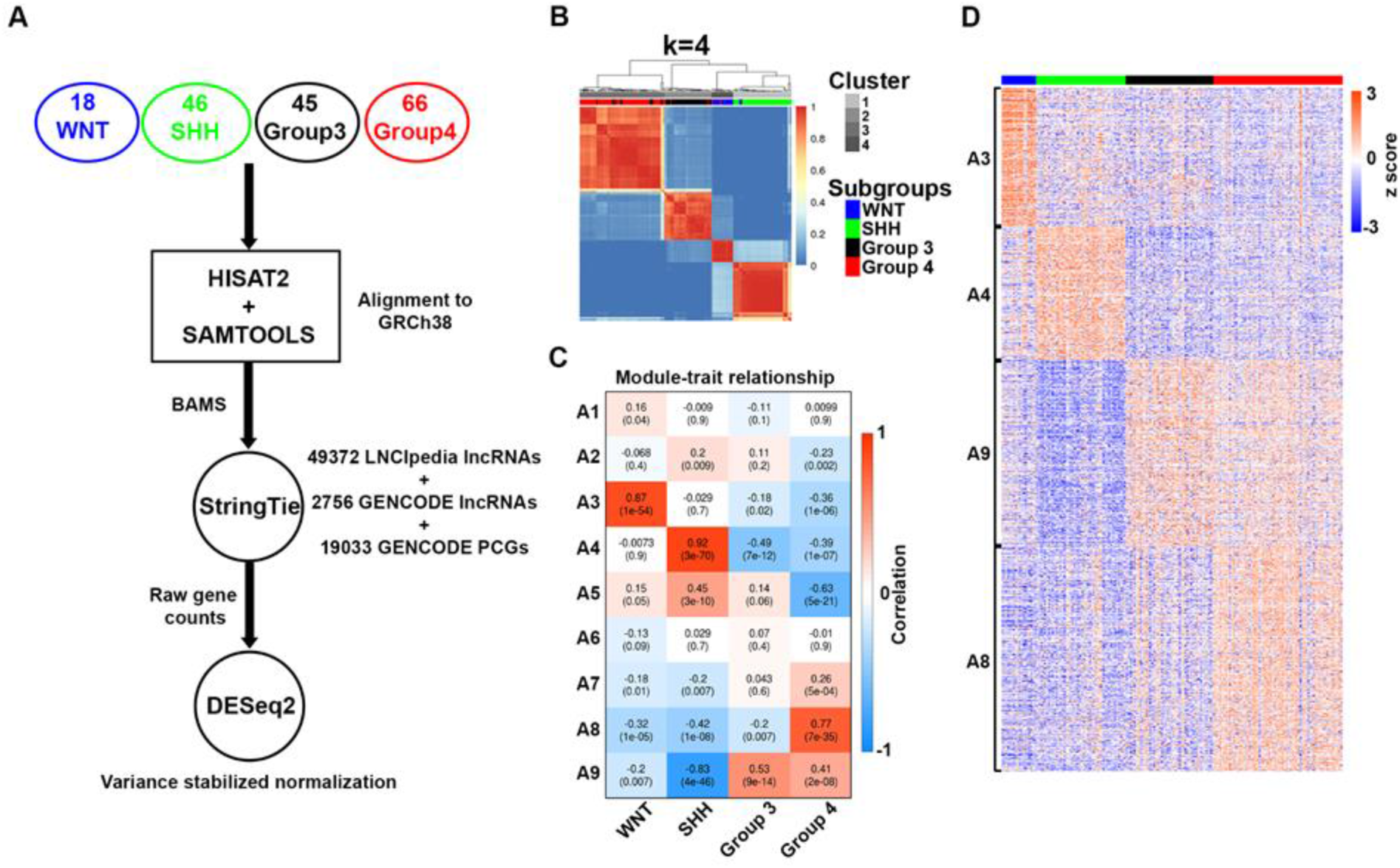
Long non-coding RNA profiles of medulloblastoma. (A) Schematic of raw data processing and analysis for medulloblastoma (MB) patients belonging to four subgroups: WNT, SHH, group 3 and group 4. (B) Heatmap showing cluster stability obtained from 1000 permutations of k-mean based clustering of 175 MB patients with top 10,000 variably expressed lncRNAs as variables. Color range depicts samples never clustered together (0, blue) to always clustered together (1, red). (C) Heatmap showing correlations of identified lncRNA cohorts (y-axis) from highly variable 5000 lncRNAs with MB subgroup phenotype (x-axis). Values in a cell show correlation level (above) and significance p-value (below in brackets). (D) Heatmap showing scaled expression level of lncRNAs in the identified cohorts (y-axis) across samples belonging to MB subgroups (x-axis).

With the objective of identifying highly variable lncRNAs specifically enriched in each subgroup, we performed weighted correlation analysis using WGCNA to find expressional co-related lncRNAs and their subgroup specificity. Weighted co-expression analysis of the top 5000 highly variable lncRNAs identified nine distinct cohorts after merging modules below the threshold. We next determined subgroup-specific module expression by performing module-trait association to obtain each module’s correlation and significance value (**Fig 1C, Table S4**). Module cohorts significantly positively correlated with WNT (467 lncRNAs, A3), SHH (452 lncRNAs, A4), group 3 (629 lncRNAs, A9), and group 4 (760 lncRNAs, A8) MBs; respectively, were identified. Gene expression in each of the identified modules also showed that these genes are highly co-expressed in their respective groups compared to other groups (**Fig 1D**). In addition, cohorts enriched in group 3 and group 4 were more correlated than WNT and SHH MBs, and vice versa. This suggests that similar to protein-coding gene expression and DNA methylation patterns, group 3 and group 4 patients also share similarities based on lncRNAs’ expression.

### A candidate diagnostic lncRNA signature for medulloblastoma subgroup classification

WGCNA analysis suggested a number of lncRNAs with subgroup-specific expression. We therefore proceeded to identify the minimum number of lncRNAs that could faithfully classify MB subgroups. To achieve our objective, we used random forest based machine learning approach that has been shown to be a robust method for such classification objectives (Mehrian-Shai et al., 2007). As patients were not evenly distributed between the four subgroups, we adopted a two-step approach: first, we identified a signature for groups with similar patient distributions i.e., SHH, group 3, and group 4; second, we identified a signature distinguishing WNT from the other subgroups (**Fig 2A**). Using this two-step approach, an 11-lncRNA signature was identified with an average <7% class error rate. Using the 11-lncRNA model, the 175 samples were re-classified into the already known subgroups with few misclassifications (**Fig 2B**), with individual lncRNA showing highly subgroup specific up/down expression (**Fig 2C**). Importantly, patient ICGC_MB23, which was classified as SHH in our random forest model but labeled WNT in the obtained dataset, was originally considered as an SHH MB in Kool et al. (Kool et al., 2014) and lacked the signature mutation in β-catenin. We also validated the 11-lncRNA based patient grouping using t-SNE based clustering (**Fig 2D**, that did not classify ICGC_MB23 as SHH) and specificity and sensitivity of the model using ROC/AUC analysis performing one versus rest (combined) comparison (**Fig 2E**).

**Figure 2.**
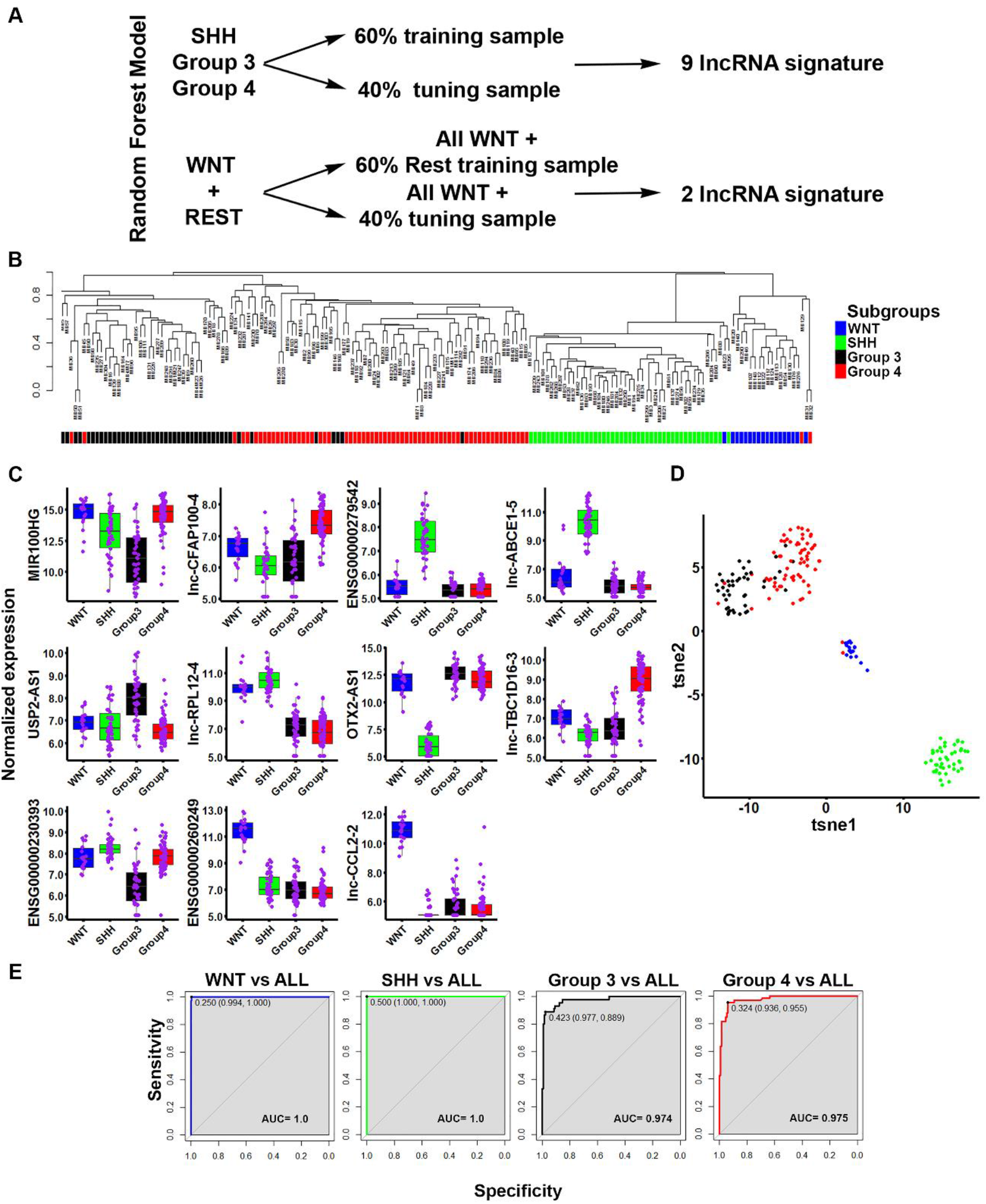
Random forest-based approach identifies an 11-lncRNA model to classify medulloblastoma subgroups. (A) Schematic depiction of the modeling process. First, a 9-lncRNA model distinguishing SHH, group 3, and group 4 patients was obtained using a 60%-40% training-tuning partition. Then, a 2-lncRNA model distinguishing WNT from the rest of the group was obtained by combining all WNT samples with a 60%-40% partition of the other subgroup patients in a training-tuning model. (B) MB patient subgroups as identified from the random forest model using only 11-lncRNA expression as variables. Dendrogram representing hierarchical clustering of dissimilarity values obtained from random forest-based classification. ICGC_MB23 is the sole WNT MB patient that clusters with the rest of the SHH MBs. Bottom color bars represent known clinical groupings (blue=WNT, green=SHH, black=group 3, red=group 4). (C) Boxplot showing distribution of normalized expression values of identified 11-lncRNAs in each patient subgroup. Purple dots represent normalized expression values for a patient. (D) tSNE plot (parameters: seed=458, perplexity=25, dims=2, eta=400, max_iter=2000) showing clustering of patients into four subgroups based on normalized expression level of identified 11-lncRNAs (blue=WNT, green=SHH, black=group 3, red=group 4). (E) ROC analysis of the linear model based on normalized expression of the identified 11-lncRNAs comparing one versus rest classifications for each of the subgroups.

In the absence of an independent dataset containing expression levels of the 11 candidate lncRNAs to validate the model, we instead validated our random forest model building process. We performed a similar classification of 175 MB samples using protein-coding genes to produce a 14-PCG model with equivalent success to the lncRNA model in classifying patient samples into the four subgroups (**Fig S2**). We then validated the 14-PCG model using the independent MAGIC microarray dataset of 763 patient belonging to the four subgroups. As expected, the 14-PCG model performed well with an overall <4% class error rate, validating our random forest model building process (**Fig S3**).

Group 3 and group 4 represent the two most heterogeneous yet closely related and difficult to distinguish MB subgroups, a pattern evident from diagnostic signature-based clustering (**Fig 2D**). To identify lncRNAs that could distinguish group 3 and group 4 tumors, we again used our random forest model approach to select highly differentially regulated and discriminative genes (**Table S5**). Our analysis yielded an 8-lncRNA model that did not improve the overall efficiency of group 3 versus group 4 classification (**Fig S4**) compared to the 11-lncRNA model distinguishing all subgroups (**Fig 2**). Nevertheless, the analysis did reveal some lncRNAs with potential functional roles in group 3 or group 4 MBs (**Fig S4B**), some of which overlapped with the 11-lncRNA model (i.e., *MIR100HG, USP2-AS1*, and *lnc-CFAP100-4*). However, we also identified other candidate lncRNAs including *ARHGEF7-AS2*, *lnc-HLX-1, lnc-EXPH5-2, lnc-CH25H-2 and lnc-TDRP-3* that showed group-specific differential expression in group 3 or group 4 patients (**Fig S4B**).

We further validated our random forest-based model in patient derived xenograft (PDX) samples derived from SHH (BT084, DMB012, RCMB32, and MED1712FH), group 3 (RCMB28, MB002, MB511H, and RCMB40), and group 4 (RCMB51, DMB006, RCMB45 and RCMB38) patients using the 9-lncRNA signature to classify SHH, group 3, and group 4 patients (**Fig 2A**), as WNT PDXs were not available for analysis. SHH, group 3, and group 4 samples were successfully identified using k-mean based clustering, principal component analysis (PCA) (**Fig 3A**) and hierarchical clustering using normalized RNA-seq expression levels (**Fig 3B**), with the exception of RCMB28 that was found to be more related to SHH PDXs. Quantitative expression of signature genes was validated by qPCR (**Fig 3C and D**) and closely resembled expression in patient RNA-seq data (**Fig 2C, Fig S5**).

**Figure 3.**
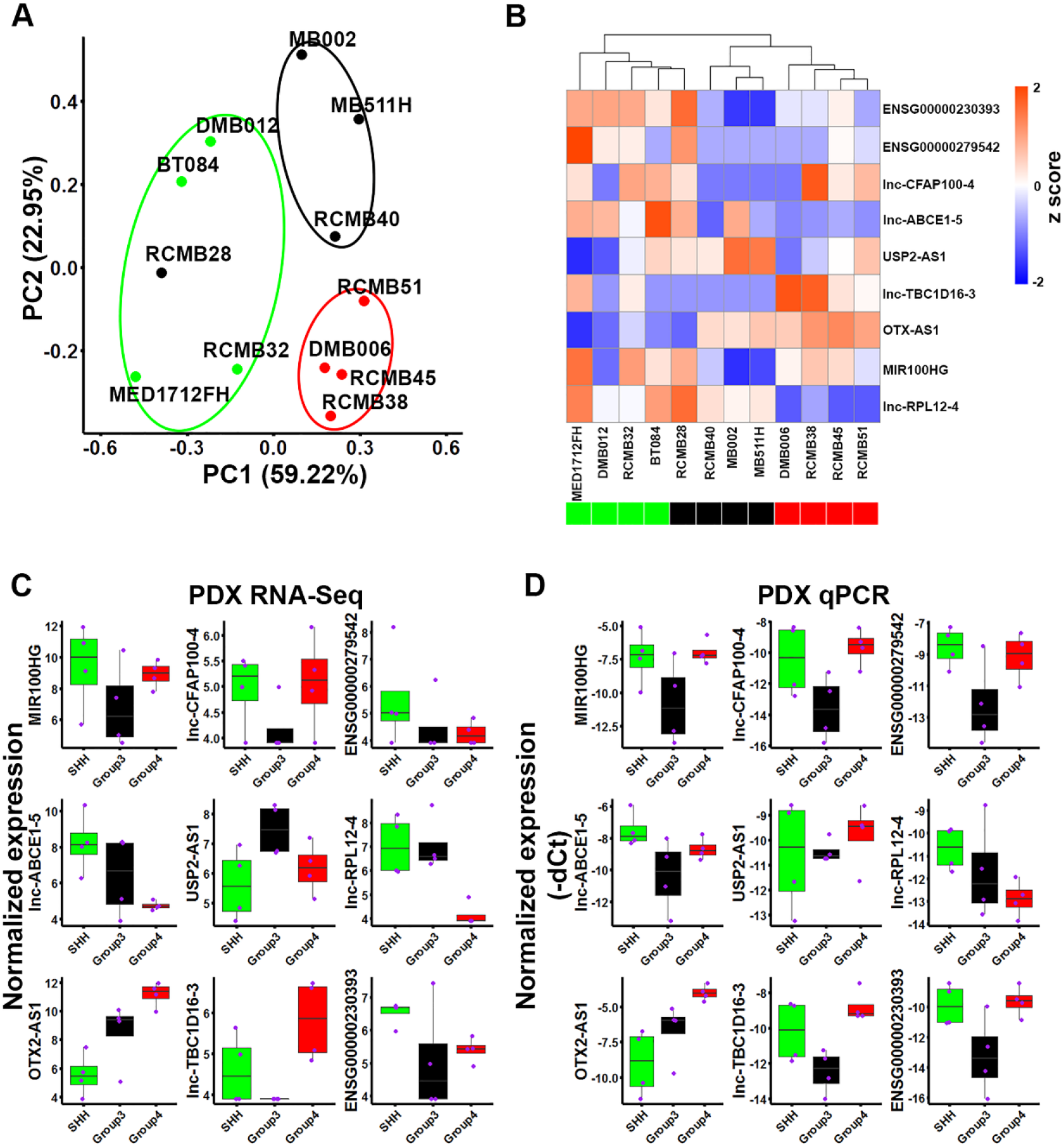
Candidate lncRNAs successfully classify PDX samples into medulloblastoma subgroups. PDX sample clustering obtained using normalized expression (RNA-seq) of 9-lncRNAs (9-lncRNA model distinguishing SHH, group 3, and group 4) in each PDX sample as the variable via (A) k-means clustering superimposed on principal component analysis (PCA), and (B) hierarchical clustering. Boxplot distributions of expression levels of the identified 9-lncRNAs from (C) RNA-seq and (D) qPCR analysis (-dCt = Ct (*candidate*) – Ct (*ACTB*)). Purple dots represent the expression level in a sample belonging to the known MB subgroup.

In order to infer the physiological importance of the identified 11-lncRNA candidates, we used a combination of CLR, arcane, and mrnet transcriptional inference algorithms to identify potential interactions between the identified lncRNAs with the expressed transcription factors in MB patients. The identified lncRNAs could potentially interact with a number of transcription factors in a complex interconnected network with both highly positively and negatively correlated associations, suggesting putative biological cross-regulation (**Fig S6**).

### Prognostic lncRNAs in medulloblastoma

To identify prognostic lncRNAs, we used the MAGIC microarray array dataset containing survival data for 612 (out of 763) patients. The microarray expression data contained expression levels of 621 lncRNAs that we used for multivariate Cox proportional hazards (Cox-PH) regression analysis. For feature selection, we utilized a penalized multivariate Cox-PH model using the LASSO penalty (α = 1). We first used the entire dataset to select 17 lncRNAs as prognostic markers and their associated penalized coefficients (**Table 1**). Of these 17 lncRNAs, 10 were markers of good prognosis and 7 were associated with poor prognosis. The 17-lncRNAs model was validated on 1000 bootstraps of the MAGIC dataset, which showed predictive stability of the proposed prognostic model. Using the penalized coefficients and log-normalized expression values for each lncRNA, we assigned a risk score to each patient. Kaplan-Meier analysis of 612 patients using the risk score as the input variable suggested that our risk score signature was a highly significant in prognostic value (HR= 13.6301, 95% CI= 8.857-20.98, logrank p-value=< 2e-16) and low score group (split at median score) represented good prognosis (Low score HR= 0.207, 95% CI= 0.133-0.323, logrank p-value= 2e-14) (**Fig 4A**). To validate our 17-lncRNA prognostic model in an independent dataset, we used ICGC RNA-seq data of 74 patients with survival information and used the penalized coefficients obtained from the MAGIC dataset analysis and variance stabilized expression from RNA-seq data to obtain an equivalent risk score. Again, Kaplan-Meier analysis showed that prognostic significance of our 17-lncRNA model (Low score HR= 0.135, 95% CI= 0.017-1.08, logrank p-value= 0.026) (**Fig 4B**). These candidate lncRNAs could potentially be involved in MB development as pro or anti-tumorigenic factors.

**Table 1.** Prognostic long non-coding RNA signature genes and associated penalized Cox-PH coefficient

|  |  |  |
| --- | --- | --- |
| **Ensembl Gene ID** | **Gene Symbol** | **Penalized coefficient** |
| ENSG00000124915 | *lnc-TMEM258-3* | -0.47771 |
| ENSG00000130600 | *H19* | 0.102589 |
| ENSG00000163009 | *lnc-RRM2-3* | 0.107783 |
| ENSG00000177993 | *ZNRF3-AS1* | -0.24098 |
| ENSG00000186960 | *LINC01551* | 0.073539 |
| ENSG00000197251 | *LINC00336* | 0.042296 |
| ENSG00000205444 | *lnc-CDYL-1* | 0.198379 |
| ENSG00000231010 | *lnc-PRR34-1* | -0.0379 |
| ENSG00000231160 | *KLF3-AS1* | -0.05371 |
| ENSG00000234665 | *lnc-FOXD4L5-25* | -0.03664 |
| ENSG00000235954 | *TTC28-AS1* | -0.00891 |
| ENSG00000244620 | *lnc-TMEM121-3* | -0.17041 |
| ENSG00000255650 | *FAM222A-AS1* | 0.007403 |
| ENSG00000256124 | *LINC01152* | -0.0675 |
| ENSG00000267278 | *MAP3K14-AS1* | -0.07358 |
| ENSG00000272841 | *AL139393.2* | 0.231787 |
| ENSG00000276399 | *AC209154.1* | -0.01405 |

**Figure 4.**
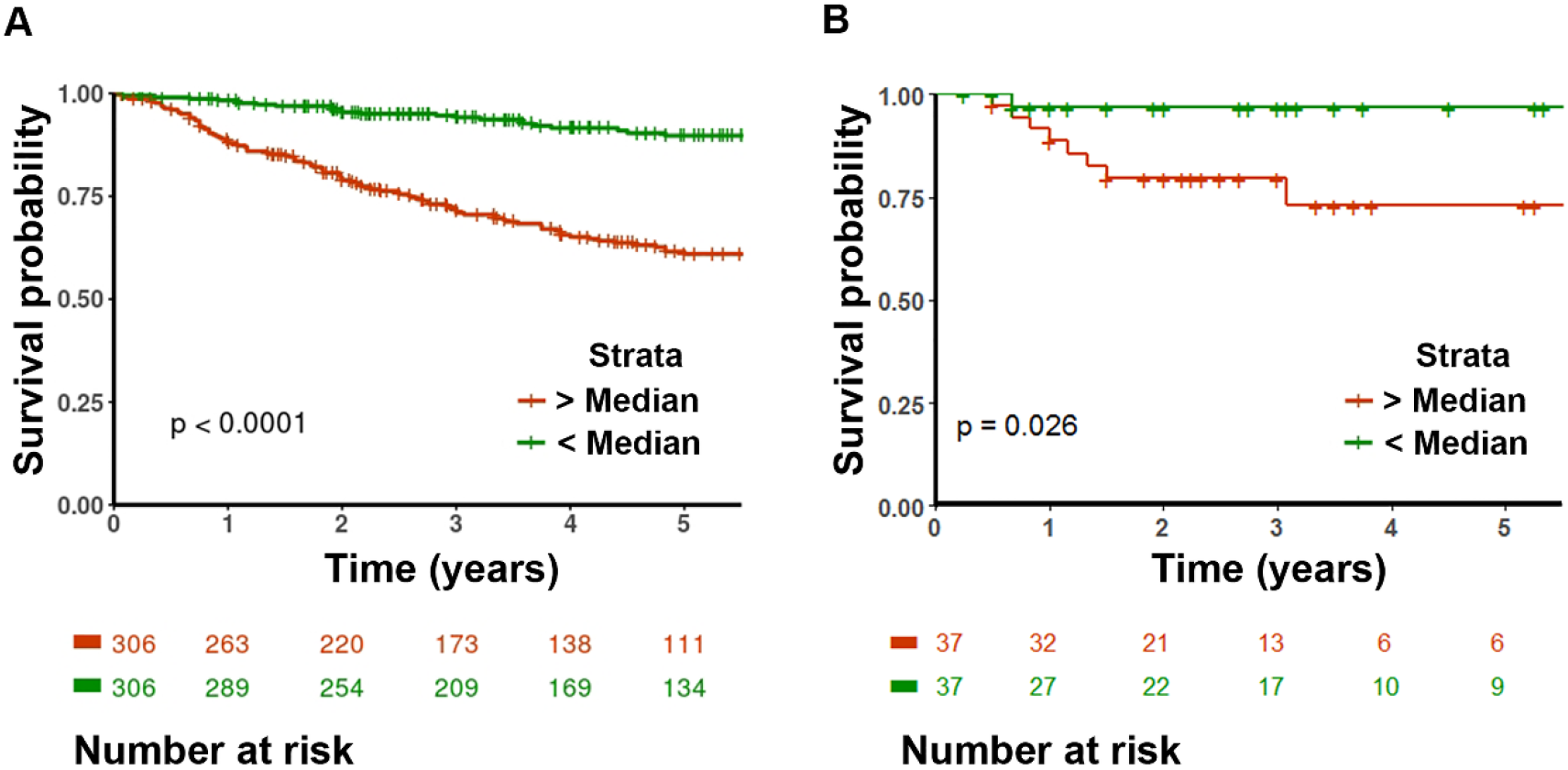
Penalized Cox proportional hazards-based lncRNA model classifies medulloblastoma patients into high and low risk groups. (A) Patients (612 MAGIC dataset) were grouped into two groups based on risk score (above and below median risk score) derived from expression of candidate prognostic lncRNAs significantly differing in their survival probability. (B) Risk score derived from the expression of the same candidate lncRNAs (except *lnc-TMEM121-3*, not detected in RNA-seq) in an independent patient cohort (74 patients in ICGC RNA-seq dataset).

## Discussion

Long non-coding RNAs are increasingly recognized as important players in cancer research (Huarte, 2015), particularly as biomarkers and/or therapeutic targets (Arriaga-Canon et al., 2018; Bhan et al., 2017; Qi and Du, 2013; Roy et al., 2018; Sarfi et al., 2019), including in brain tumors (Li et al., 2018; Liang et al., 2019; Pop et al., 2018; Reon et al., 2016; Zeng et al., 2018). However, there is a lack of knowledge of lncRNAs’ involvement in MBs. Here we bridged this knowledge gap by proposing diagnostic and prognostic biomarkers candidates for further study *in vitro* and *in vivo* systems to understand their potential function in MB genesis/progression. Our study is the first genome-wide analysis of lncRNAs’ expression profile in MB and its subgroups. Overall lncRNAs’ expression dynamics mirrors the well-known MB heterogeneity seen in genetic and epigenetic analyses (Cavalli et al., 2017; Northcott et al., 2017). MB subgroup clusters obtained using highly variable lncRNAs overlapped with existing clinical and molecular subgroups. Using variantly expressed lncRNAs and weighted correlations, we further identified subgroup-specific lncRNAs. These upregulated lncRNAs might represent functionally relevant genes and require further validation. The obtained lncRNA signature could be curated using transcriptional inference algorithms and proximity to or co-relation with known MB relevant protein coding genes for further functional validation *in vitro* and *in vivo* studies.

Presently, very few lncRNAs have been studied for their putative roles in MB or its subtypes. *NKX2-2AS* was shown *in vitro* to modulate SHH-potentiated MB development by acting as a miRNA sponge for miR-103 and miR-107, thereby de-repressing their tumor suppressive targets BTG2 and LATS1 and inhibiting proliferation and migration (Zhang et al., 2018). *CDKN2B-AS1 (ANRIL)* has been shown to promote proliferation *in vitro* studies by sponging miR-323 and activating BRI3 dependent p38-MAPK, AKT and WNT signaling (Zhang et al., 2017), and in current analysis, it was found to be upregulated in group 4 patients compared to other MBs. *PVT1* is prevalently found fused to MYC and NDRG1 genes in group 3 tumors, leading to oncogenic transformation of these genes (Northcott et al., 2012). *lnc-IRX3-80 (CRNDE)* was also reported as an oncogenic lncRNA *in vitro* and *in vivo* studies (Song et al., 2016). Both *PVT1* and *lnc-IRX3-80* were upregulated in WNT and SHH MBs in our current analysis*. Lnc-FAM84B-15 (CCAT1)*, which was found upregulated in WNT and group 3 MBs, has also been shown to be involved in promoting tumor proliferation and metastasis by activating MAPK pathway (Gao et al., 2018). *MIR100HG (lnc-NeD125)* has been shown to be overexpressed in group 4 MBs, again acting as an miRNA sponge for miR-19a-3p, miR-19b-3p and miR-106a-5p, exerting an oncogenic function by de-repressing cell cycle target genes (Laneve et al., 2017). *MIR100HG* is also oncogenic in gastric cancer (Li et al., 2019), breast cancer (Wang et al., 2018b), and leukemia (Emmrich et al., 2014). In our analysis, out of the above described lncRNAs only *MIR100HG (lnc-NeD125)* was selected in our diagnostic signature, being highly expressed in all MBs but group 3 (**Fig 2**). The rest of the diagnostic markers have not yet been studies in medulloblastoma or other tumors to our current knowledge. However, our 11-lncRNAs model could complement existing molecular and clinical-based diagnostic approaches, particularly for group 3 and group 4 MBs. Some of the identified signature lncRNAs are highly subgroup-specific, such as: *lnc-CCL2-2* (WNT), *lnc-ABCE1-5* (SHH), *USP2-AS1* (group 3), and *lnc-TBC1D16-3* (group 4). Our mutual information-based network analysis also identified putative interacting transcription factors involved in medulloblastoma and other cancers (**Fig S6**); for example, FOXO1 (Pei et al., 2016; Srivastava et al., 2009), OTX2 (Lu et al., 2017), NRL, CRX (Garancher et al., 2018) and TET3 (Bezerra Salomao et al., 2018). These predicted interactions could be tested for their relevance in functional role of identified biomarker candidates in medulloblastoma pathogenesis in future follow up studies.

Our 17-lncRNAs prognostic model represents another set of putative functionally important lncRNAs. Of the 17 lncRNAs, seven were associated with poor prognosis, including *H19 (***Table 1**). None of the candidate prognostic markers (poor prognosis) were enriched (highly expressed) in a specific subgroup of patients (**Fig S7A, B**). Some candidate poor prognostic markers were positively correlated with worse prognosis MBs subgroups (e.g.:*H19* expression was positively correlated with SHH MBs associated diagnostic lncRNA candidates, **Fig S6 C**) as expected from poor survivability of those subgroup patients. However, none of the prognostic markers were highly correlated (correlation >0.80) with any 11 diagnostic candidate lncRNA. Hence, these prognostic markers could represent a set of subgroup independent lncRNA candidates that drive MB progression. *H19* is a well-studied oncogenic lncRNA in various cancer systems including glioblastoma, where it has been shown to be promote cellular proliferation and metastasis (Fazi et al., 2018; Jiang et al., 2016; Pei et al., 2016; Raveh et al., 2015; Zhou et al., 2017). LncRNA *LINC01551* has been found to upregulate cellular proliferation and migration in non-CNS cancers such as hepatocellular carcinoma by interacting with the miR122-ADAM10 axis (Gao et al., 2019). *LINC00336* promoted lung cancer progression by inhibiting regulated cell death by knocking down miR-6852 function (Wang et al., 2019).

Overall, our analysis proposes new lncRNAs candidates in MB with functional, diagnostic, and prognostic significance that warrant further investigation and validation. This is the first global analysis of lncRNAs in MB that will provide an invaluable resource for those working in the field to prioritize for further study.

## Data Availability

The 175 MB patients’ RNA-seq data is available from the European Genome-Phenome Archive (EGA, http://www.ebi.ac.uk/ega/), accession number EGAD00001003279 (Northcott et al., 2017). Microarray expression datasets from 763 medulloblastoma patients is available at Gene Expression Omnibus Accession Number GSE85218 (Cavalli et al., 2017). RNA-seq for PDX tissues generated in the lab is available at Gene Expression Omnibus Accession Number GSE134248.

## Acknowledgements

We are thankful to Dr. Alexandra Garancher and Dr. Robert Wechsler-Reya (SBP, San Diego) for sharing MB PDX tissue pellets. This work was supported by National Institutes of Health grants NCI 5P30CA030199 (SBP), P30 CA006973 (JHU SKCCC) and Florida Department of Health, Bankhead-Coley Cancer Research Program 5BC08 to R.J.P.

## Competing interests

The authors declare that they have no competing interests.

## Notes

#### Summary of Updates

Made revision as per reviewers' comments.

## References

Arriaga-Canon, C., et al., 2018. The use of long non-coding RNAs as prognostic biomarkers and therapeutic targets in prostate cancer. Oncotarget. 9, 20872–20890.

Bezerra Salomao, K., et al., 2018. Reduced hydroxymethylation characterizes medulloblastoma while TET and IDH genes are differentially expressed within molecular subgroups. J Neurooncol. 139, 33–42.

Bhan, A., et al., 2017. Long Noncoding RNA and Cancer: A New Paradigm. Cancer Res. 77, 3965–3981.

Cavalli, F. M. G., et al., 2017. Intertumoral Heterogeneity within Medulloblastoma Subgroups. Cancer Cell. 31, 737–754.e6.

De Braganca, K. C., Packer, R. J., 2013. Treatment Options for Medulloblastoma and CNS Primitive Neuroectodermal Tumor (PNET). Curr Treat Options Neurol. 15, 593–606.

Diamandis, P., Aldape, K., 2018. World Health Organization 2016 Classification of Central Nervous System Tumors. Neurol Clin. 36, 439–447.

Dufour, C., et al., 2012. Metastatic Medulloblastoma in Childhood: Chang’s Classification Revisited. Int J Surg Oncol. 2012, 245385.

Emmrich, S., et al., 2014. LincRNAs MONC and MIR100HG act as oncogenes in acute megakaryoblastic leukemia. Mol Cancer. 13, 171.

Fazi, B., et al., 2018. The lncRNA H19 positively affects the tumorigenic properties of glioblastoma cells and contributes to NKD1 repression through the recruitment of EZH2 on its promoter. Oncotarget. 9, 15512–15525.

Frankish, A., et al., 2019. GENCODE reference annotation for the human and mouse genomes. Nucleic Acids Res. 47, D766–d773.

Gao, J., et al., 2019. Long noncoding LINC01551 promotes hepatocellular carcinoma cell proliferation, migration, and invasion by acting as a competing endogenous RNA of microRNA-122-5p to regulate ADAM10 expression. J Cell Biochem.

Gao, R., et al., 2018. Long noncoding RNA CCAT1 promotes cell proliferation and metastasis in human medulloblastoma via MAPK pathway. Tumori. 104, 43–50.

Garancher, A., et al., 2018. NRL and CRX Define Photoreceptor Identity and Reveal Subgroup-Specific Dependencies in Medulloblastoma. Cancer Cell. 33, 435–449.e6.

Huarte, M., 2015. The emerging role of lncRNAs in cancer. Nat Med. 21, 1253–61.

Iyer, M. K., et al., 2015. The landscape of long noncoding RNAs in the human transcriptome. Nat Genet. 47, 199–208.

Jiang, X., et al., 2016. Increased level of H19 long noncoding RNA promotes invasion, angiogenesis, and stemness of glioblastoma cells. J Neurosurg. 124, 129–36.

Jones, D. T., et al., 2012. Dissecting the genomic complexity underlying medulloblastoma. Nature. 488, 100–5.

Kondoff, S. I., et al., 2015. A case of early extraneural medulloblastoma metastases in a young adult. Asian J Neurosurg. 10, 331–3.

Kool, M., et al., 2014. Genome sequencing of SHH medulloblastoma predicts genotype-related response to smoothened inhibition. Cancer Cell. 25, 393–405.

Laneve, P., et al., 2017. The long noncoding RNA linc-NeD125 controls the expression of medulloblastoma driver genes by microRNA sponge activity. Oncotarget. 8, 31003–31015.

Langfelder, P., Horvath, S., 2008. WGCNA: an R package for weighted correlation network analysis. BMC Bioinformatics. 9, 559.

Li, J., et al., 2019. MIR100HG: a credible prognostic biomarker and an oncogenic lncRNA in gastric cancer. Biosci Rep. 39.

Li, J., et al., 2018. Targeting Long Noncoding RNA in Glioma: A Pathway Perspective. Mol Ther Nucleic Acids. 13, 431–441.

Liang, R., et al., 2019. Analysis of long non-coding RNAs in glioblastoma for prognosis prediction using weighted gene co-expression network analysis, Cox regression, and L1-LASSO penalization. Onco Targets Ther. 12, 157–168.

Lin, C. Y., et al., 2016. Active medulloblastoma enhancers reveal subgroup-specific cellular origins. Nature. 530, 57–62.

Long, Y., et al., 2017. How do lncRNAs regulate transcription? Sci Adv. 3, eaao2110.

Louis, D. N., et al., 2007. The 2007 WHO classification of tumours of the central nervous system. Acta Neuropathol. 114, 97–109.

Love, M. I., et al., 2014. Moderated estimation of fold change and dispersion for RNA-seq data with DESeq2. Genome Biol. 15, 550.

Lu, Y., et al., 2017. OTX2 expression contributes to proliferation and progression in Myc-amplified medulloblastoma. Am J Cancer Res. 7, 647–656.

Martin, A. M., et al., 2014. Management of pediatric and adult patients with medulloblastoma. Curr Treat Options Oncol. 15, 581–94.

Mehrian-Shai, R., et al., 2007. Insulin growth factor-binding protein 2 is a candidate biomarker for PTEN status and PI3K/Akt pathway activation in glioblastoma and prostate cancer. Proc Natl Acad Sci U S A. 104, 5563–8.

Meyer, P. E., et al., 2008. minet: A R/Bioconductor package for inferring large transcriptional networks using mutual information. BMC Bioinformatics. 9, 461.

Millard, N. E., De Braganca, K. C., 2016. Medulloblastoma. J Child Neurol. 31, 1341–53.

Northcott, P. A., et al., 2017. The whole-genome landscape of medulloblastoma subtypes. Nature. 547, 311–317.

Northcott, P. A., et al., 2014. Enhancer hijacking activates GFI1 family oncogenes in medulloblastoma. Nature. 511, 428–34.

Northcott, P. A., et al., 2019. Medulloblastoma. Nat Rev Dis Primers. 5, 11.

Northcott, P. A., et al., 2012. Subgroup-specific structural variation across 1,000 medulloblastoma genomes. Nature. 488, 49–56.

Ostrom, Q. T., et al., 2017. CBTRUS Statistical Report: Primary brain and other central nervous system tumors diagnosed in the United States in 2010-2014. Neuro Oncol. 19, v1–v88.

Palmer, S. L., et al., 2007. Understanding the cognitive impact on children who are treated for medulloblastoma. J Pediatr Psychol. 32, 1040–9.

Pei, Y., et al., 2016. HDAC and PI3K Antagonists Cooperate to Inhibit Growth of MYC-Driven Medulloblastoma. Cancer Cell. 29, 311–323.

Pop, S., et al., 2018. Long non-coding RNAs in brain tumours: Focus on recent epigenetic findings in glioma. J Cell Mol Med. 22, 4597–4610.

Qi, P., Du, X., 2013. The long non-coding RNAs, a new cancer diagnostic and therapeutic gold mine. Mod Pathol. 26, 155–65.

Raveh, E., et al., 2015. The H19 Long non-coding RNA in cancer initiation, progression and metastasis - a proposed unifying theory. Mol Cancer. 14, 184.

Reon, B. J., et al., 2016. Expression of lncRNAs in Low-Grade Gliomas and Glioblastoma Multiforme: An In Silico Analysis. PLoS Med. 13, e1002192.

Roy, S., et al., 2018. A General Overview on Non-coding RNA-Based Diagnostic and Therapeutic Approaches for Liver Diseases. Front Pharmacol. 9, 805.

Sahakyan, A., et al., 2018. The Role of Xist in X-Chromosome Dosage Compensation. Trends Cell Biol. 28, 999–1013.

Sarfi, M., et al., 2019. Long noncoding RNAs biomarker-based cancer assessment. J Cell Physiol.

Shan, Y., et al., 2018. LncRNA SNHG7 sponges miR-216b to promote proliferation and liver metastasis of colorectal cancer through upregulating GALNT1. Cell Death Dis. 9, 722.

Smee, R. I., et al., 2012. Medulloblastoma: progress over time. J Med Imaging Radiat Oncol. 56, 227–34.

Song, H., et al., 2016. Long non-coding RNA CRNDE promotes tumor growth in medulloblastoma. Eur Rev Med Pharmacol Sci. 20, 2588–97.

Srivastava, V. K., et al., 2009. Impaired medulloblastoma cell survival following activation of the FOXO1 transcription factor. Int J Oncol. 35, 1045–51.

Tibshirani, R., 1997. The lasso method for variable selection in the cox model. Statistics in Medicine. 16, 385–395.

Trimarchi, T., et al., 2014. Genome-wide mapping and characterization of Notch-regulated long noncoding RNAs in acute leukemia. Cell. 158, 593–606.

Uszczynska-Ratajczak, B., et al., 2018. Towards a complete map of the human long non-coding RNA transcriptome. Nat Rev Genet. 19, 535–548.

Vladoiu, M. C., et al., 2019. Childhood cerebellar tumours mirror conserved fetal transcriptional programs. Nature. 572, 67–73.

Volders, P. J., et al., 2019. LNCipedia 5: towards a reference set of human long non-coding RNAs. Nucleic Acids Res. 47, D135–d139.

Wang, J., et al., 2018a. Medulloblastoma: From Molecular Subgroups to Molecular Targeted Therapies. Annu Rev Neurosci. 41, 207–232.

Wang, M., et al., 2019. Long noncoding RNA LINC00336 inhibits ferroptosis in lung cancer by functioning as a competing endogenous RNA. Cell Death Differ.

Wang, S., et al., 2018b. LncRNA MIR100HG promotes cell proliferation in triple-negative breast cancer through triplex formation with p27 loci. Cell Death Dis. 9, 805.

Wilkerson, M. D., Hayes, D. N., 2010. ConsensusClusterPlus: a class discovery tool with confidence assessments and item tracking. Bioinformatics. 26, 1572–3.

Zeng, T., et al., 2018. Exploring Long Noncoding RNAs in Glioblastoma: Regulatory Mechanisms and Clinical Potentials. Int J Genomics. 2018, 2895958.

Zhang, H., et al., 2017. Potential Role of Long Non-Coding RNA ANRIL in Pediatric Medulloblastoma Through Promotion on Proliferation and Migration by Targeting miR-323. J Cell Biochem. 118, 4735–4744.

Zhang, X., et al., 2012. Long non-coding RNA expression profiles predict clinical phenotypes in glioma. Neurobiol Dis. 48, 1–8.

Zhang, Y., et al., 2018. Nkx2-2as Suppression Contributes to the Pathogenesis of Sonic Hedgehog Medulloblastoma. Cancer Res. 78, 962–973.

Zhou, W., et al., 2017. The lncRNA H19 mediates breast cancer cell plasticity during EMT and MET plasticity by differentially sponging miR-200b/c and let-7b. Sci Signal. 10.

