## Supplemental Information for "In silico analysis of long non-coding RNAs in medulloblastoma and its subgroups"

### Supplementary data

### Supplementary Tables

**Table S1.** List of 175 medulloblastoma patient samples and clinical information.

**Table S2.** List of primers used for qPCR validation of 9 diagnostic candidates in PDX samples.

**Table S3.** Confusion matrices associated with individual subgroups as obtained from cluster obtained with k=4.

**Table S4.** List of subgroup specific lncRNA cohorts identified by WGCNA analysis.

**Table S5.** List of differentially expressed lncRNA between group 3 and group 4 patients.

### Supplementary Figures

**Fig S1. Medulloblastoma patients can be optimally subgrouped into 4 clusters based on long non-coding RNA expression.** (A) Heatmap depicting stability of k-means clusters, 2 to 6 (k=4, in Fig 1C), based on consensus clustering. Color range depicts samples never clustered together (0, blue) to always clustered together (1, red). (B) Colored line depicting relationship between cumulative distribution function (CDF) and consensus index for each of the k-means values 2 to 6. (C) Graph showing relative change in area under the CDF curve (in B) comparing k to k-1 for k from 3 to 6. For k=2, the value is total area under the CDF curve in B. (D) Cluster consensus plot showing mean of pairwise consensus value for all cluster members. For k=4, the graph shows that each of the obtained clusters are of similar stability.

**Fig S2. Random forest-based approach identifies a 14-PCG model to classify medulloblastoma subgroups.** (A) Schematic depiction of the modeling process. First, an 11-PCG model distinguishing SHH, group 3, and group 4 patients was obtained using a 60%-40% training-tuning partition. Then, a 3-PCG model distinguishing WNT from the rest of the group was obtained by combining all WNT samples with a 60%-40% partition of the other subgroup patients in a training-tuning model. (B) MB patient subgroups as identified from the random forest model using only 14-PCG expression as variables. Dendrogram represents hierarchical clustering of dissimilarity values obtained from random forest-based classification. Bottom color bars represent known clinical groupings (blue=WNT, green=SHH, black=group 3, red=group 4). The obtained clustering and misclassification are similar to that in Fig 2B. (C) Boxplot showing distribution of normalized expression values of identified 14-PCGs in each patient subgroup. Purple dots represent normalized expression value for a patient. D) tSNE plot (parameters: seed=458, perplexity=25,dims=2,eta=400, max_iter=2000) showing clustering of patients into four subgroups based on normalized expression level of identified 14-PCGs (blue=WNT, green=SHH, black=group 3, red=group 4). The obtained clustering is similar to that in Fig 3D, except for sample ICGC_MB23 that cluster with SHH subgroup. (E) ROC analysis of linear model based on normalized expression of identified 14-PCGs comparing one versus rest classifications for each of the subgroups. The AUC values for 11-lncRNAs model and 14-PCGs model are comparable.

**Fig S3. 14-PCG model classifies independent medulloblastoma patient samples with high efficiency.** (A) MB patient subgroups as identified in the Cavalli17 dataset on applying a random forest model using only 14-PCG expression as variables. Dendrogram represents hierarchical clustering of dissimilarity values obtained from random forest-based classification. Bottom color bars represent known clinical grouping (blue=WNT, green=SHH, black=group 3, red=group 4). (B) Boxplot showing distribution of normalized expression values of identified 14-PCGs in each patient subgroup. Purple dots represent normalized expression value for a patient. The normalized expression distribution for the 14-PCGs are similar to that in RNA-seq analysis (Fig S2C). (D) tSNE plot (parameters: seed=999, perplexity=25,dims=2,eta=400, max_iter=2000) showing clustering of patients into four subgroup based on normalized expression level of identified 14-PCGs (blue=WNT, green=SHH, black=group 3, red=group 4). (D) ROC analysis of linear model based on normalized expression of identified 14-PCGs comparing one versus rest classifications for each of the subgroups.

**Fig S4. Random forest-based approach identifies an 8-lncRNA model to classify group 3 and group 4 patents.** (A) MB patient subgroups as identified from a random forest model using 8-lncRNA expression as variables. Dendrogram represents hierarchical clustering of dissimilarity values obtained from random forest-based classification. Bottom color bars represent known clinical groupings (black=group 3, red=group 4). (B) Boxplot showing distribution of normalized expression values of identified 8-lncRNAs in group 3 and group 4 MBs. Purple dots represent normalized expression value for a patient. (D) tSNE plot (parameters: seed=458, perplexity=25,dims=2,eta=400, max_iter=2000) showing clustering of group 3 and group 4 patients into two heterogeneous clusters based on normalized expression levels of identified 8-lncRNAs (black=group 3, red=group 4). (E) ROC analysis of linear model based on normalized expression of identified 8-lncRNAs comparing group 3 versus group 4 samples.

**Fig S5. Correlation between RNA-seq and qPCR data obtained from PDX samples.** Scatter plot showing positive correlation between normalized RNA-seq and qPCR values of each from all contributing 12 samples. The best-fit line represents the predicted linear model (y~x) for each plot.

**Fig S6. Putative interaction network of identified 11-lncRNAs with transcription factors.** Interaction network between the 11-lncRNAs and transcription factors based on a consensus of mutual information analysis using CLR, arcane, and mrnet-based approaches.

**Fig S7. Expression distribution of identified bad prognostic markers**. Box plot representing expression distribution of candidate bad prognostic markers in (A) MAGIC microarray dataset and (B) ICGC RNA-seq dataset in each subgroup. None of the identified lncRNAs is expressed specifically in a subgroup; however, patients expressing high levels were primarily in group 3. (C) Pearson correlation between 16 lncRNA prognostic candidates (*lnc-TMEM121-3* not detected in RNA-seq) and 11 lncRNA diagnostic candidates (marked with *). Poor prognostic markers didn’t highly correlated (>0.80) with any 11 diagnostic candidate lncRNA in particular.

**Figure S1**

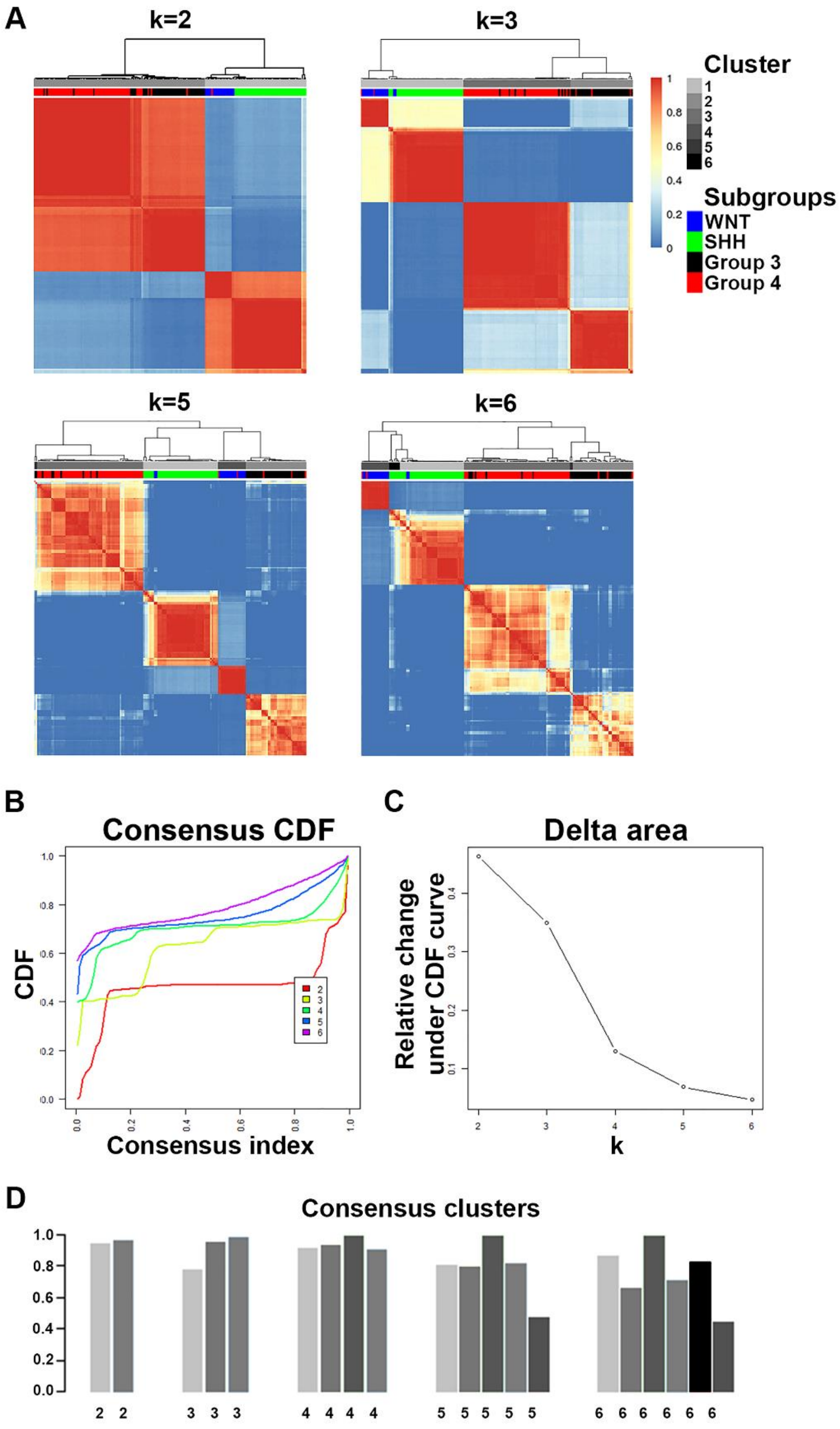

A
k=2
k=3
Cluster
1
2
3
4
5
6
Subgroups
WNT
SHH
Group 3
Group 4
k=5
k=6
B
Consensus CDF
CDF
Consensus index
C
Delta area
Relative change under CDF curve
k
D
Consensus clusters

**Figure S2**

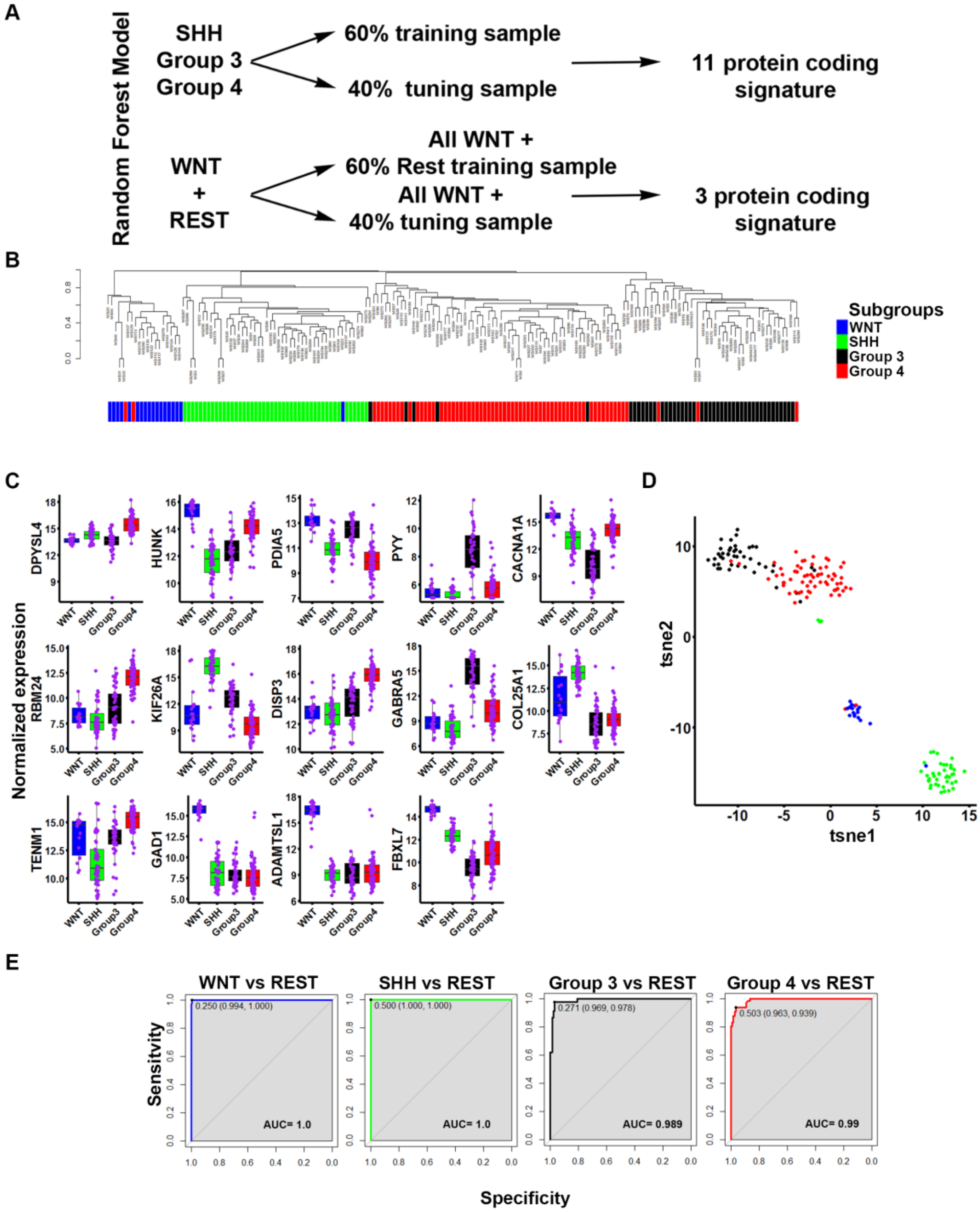

A
Random Forest Model
SHH
Group 3
Group 4
60% training sample
40% tuning sample
11 protein coding signature
WNT
+
REST
All WNT +
60% Rest training sample
All WNT +
40% tuning sample
3 protein coding signature
B
Subgroups
WNT
SHH
Group 3
Group 4
C
Normalized expression
DPYSL4
HUNK
PDIA5
PYY
CACNA1A
RBM24
KIF26A
DISP3
GABRA5
COL25A1
TENM1
GAD1
ADAMTSL1
FBXL7
WNT
SHH
Group3
Group4
D
tsne2
tsne1
E
WNT vs REST
SHH vs REST
Group 3 vs REST
Group 4 vs REST
0.250 (0.994, 1.000)
0.500 (1.000, 1.000)
0.271 (0.969, 0.978)
0.503 (0.963, 0.939)
AUC= 1.0
AUC= 1.0
AUC= 0.989
AUC= 0.99
Sensitivity
Specificity

**Figure S3**

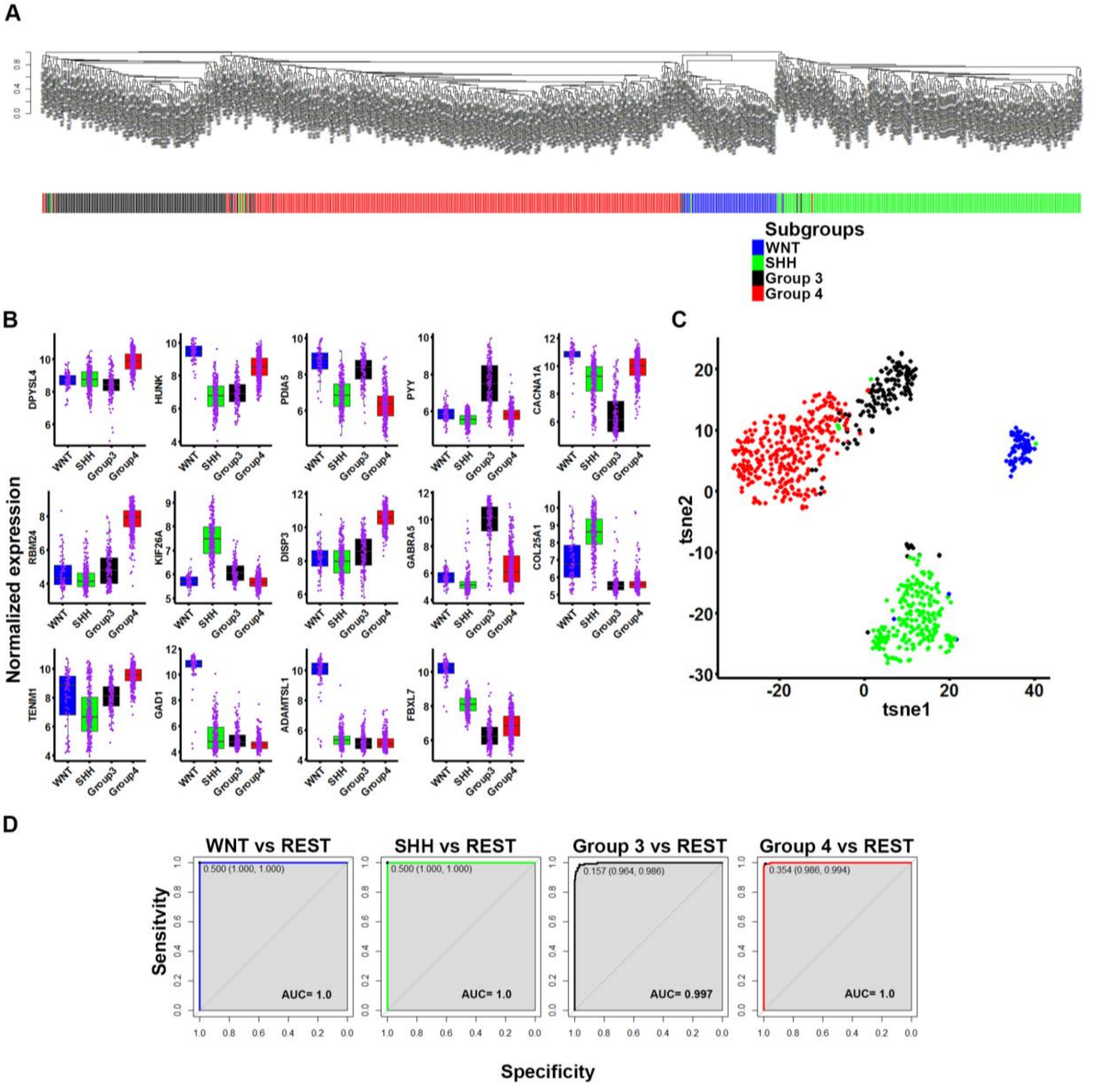
A
Subgroups
WNT
SHH
Group 3
Group 4
B
Normalized expression
DPYSL4
HUNK
PDIA5
PYY
CACNA1A
RBM24
KIF26A
DISP3
GABRA5
COL25A1
TENM1
GAD1
ADAMTSL1
FBXL7
WNT
SHH
Group3
Group4
C
tsne2
tsne1
D
WNT vs REST
SHH vs REST
Group 3 vs REST
Group 4 vs REST
0.500 (1.000, 1.000)
0.500 (1.000, 1.000)
0.157 (0.964, 0.986)
0.354 (0.986, 0.994)
AUC= 1.0
AUC= 1.0
AUC= 0.997
AUC= 1.0
Sensitivity
Specificity

**Figure S4**

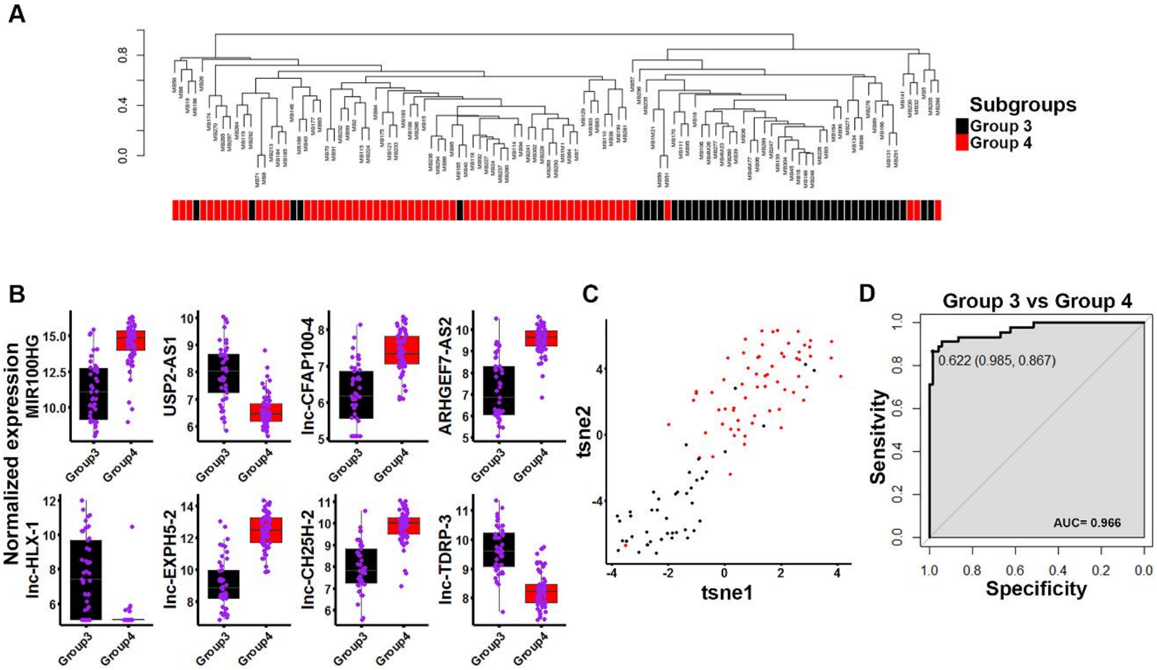

A
Subgroups
Group 3
Group 4
B
Normalized expression
MIR100HG
USP2-AS1
lnc-CFAP100-4
ARHGEF7-AS2
lnc-HLX-1
lnc-EXPH5-2
lnc-CH25H-2
lnc-TDRP-3
Group3
Group4
C
tsne1
tsne2
D
Group 3 vs Group 4
0.622 (0.985, 0.867)
AUC= 0.966
Sensitivity
Specificity

**Figure S5**

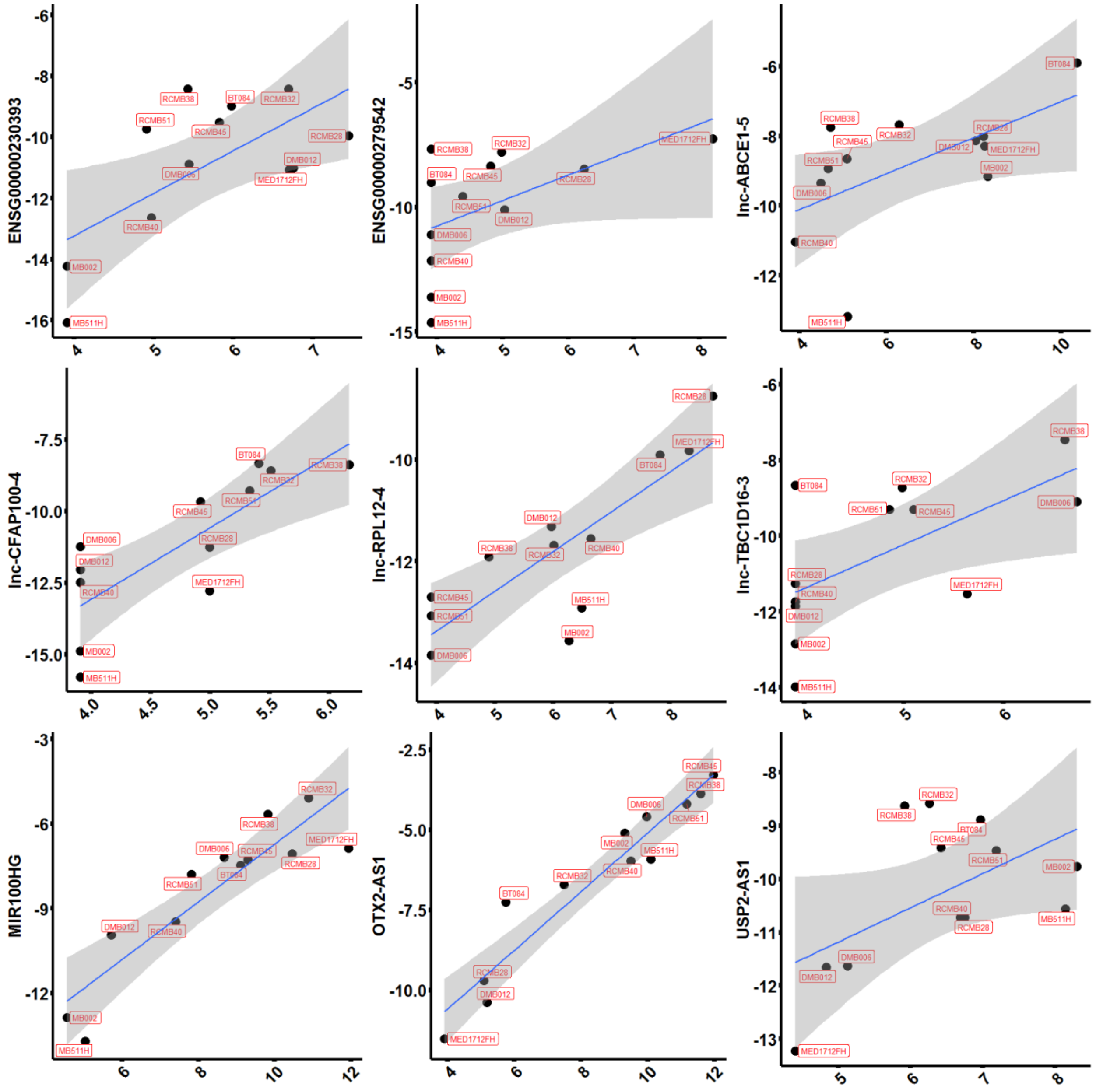

ENSG00000230393
ENSG00000279542
lnc-ABCE1-5
lnc-CFAP100-4
lnc-RPL12-4
lnc-TBC1D16-3
MIR100HG
OTX2-AS1
USP2-AS1

**Figure S6**

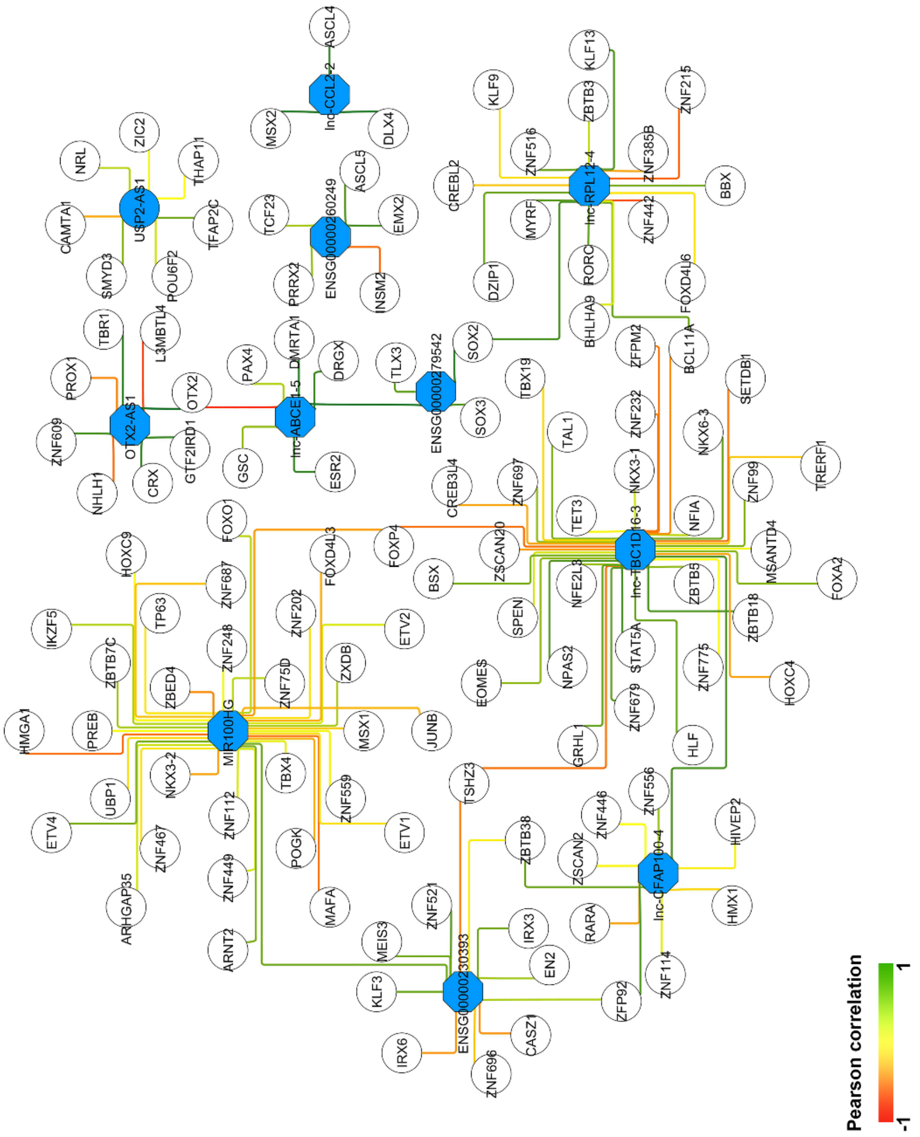

USP2-AS1
OTX2-AS1
lnc-ABCE1-5
lnc-CCL22-2
ENSG00000260249
ENSG00000279542
lnc-RPL12-4
lnc-TBC1D16-3
MIR100HG
ENSG00000230393
lnc-CFAP100-4
NRL
ZIC2
THAP11
TFAP2C
CAMTA1
SMYD3
POU6F2
TBR1
L3MBTL4
PROX1
OTX2
ZNF609
NHLH1
CRX
GTF2IRD1
PAX4
DMRTA1
DRGX
GSC
ESR2
MSX2
ASCL4
DLX4
TCF23
ASCL5
EMX2
PRRX2
INSM2
TLX3
SOX2
SOX3
KLF9
ZNF516
ZBTB3
KLF13
ZNF215
CREBL2
MYRF
ZNF385B
BBX
ZNF442
DZIP1
RORC
FOXD4L6
BHLHA9
ZFPM2
BCL11A
TBX19
ZNF232
SETDB1
TAL1
NKX6-3
TRERF1
CREB3L4
ZNF697
NKX3-1
ZNF99
TET3
NFIA
MSANTD4
FOXA2
FOXO1
FOXD4L3
FOXP4
ZSCAN20
HOXC9
ZNF687
BSX
NFE2L3
ZBTB5
ZBTB18
IKZF5
TP63
ZNF202
ETV2
SPENI
ZBTB7C
ZNF248
ZXDB
NPAS2
STAT5A
ZNF775
HOXC4
ZBED4
ZNF75D
EOMES
ZNF679
HMGA1
PREB
MSX1
JUNB
GRHL1
HLF
UBP1
NKX3-2
TBX4
TSHZ3
ZNF556
ETV4
ZNF112
ZNF559
ZNF446
HIVEP2
ZNF467
POGK
ETV1
ZBTB38
ZSCAN2
ARHGAP35
ZNF449
MAFA
ZNF521
IRX3
RARA
HMX1
ARNT2
MEIS3
EN2
ZNF114
KLF3
ZFP92
IRX6
CASZ1
ZNF696
Pearson correlation
-1
1

**Figure S7**

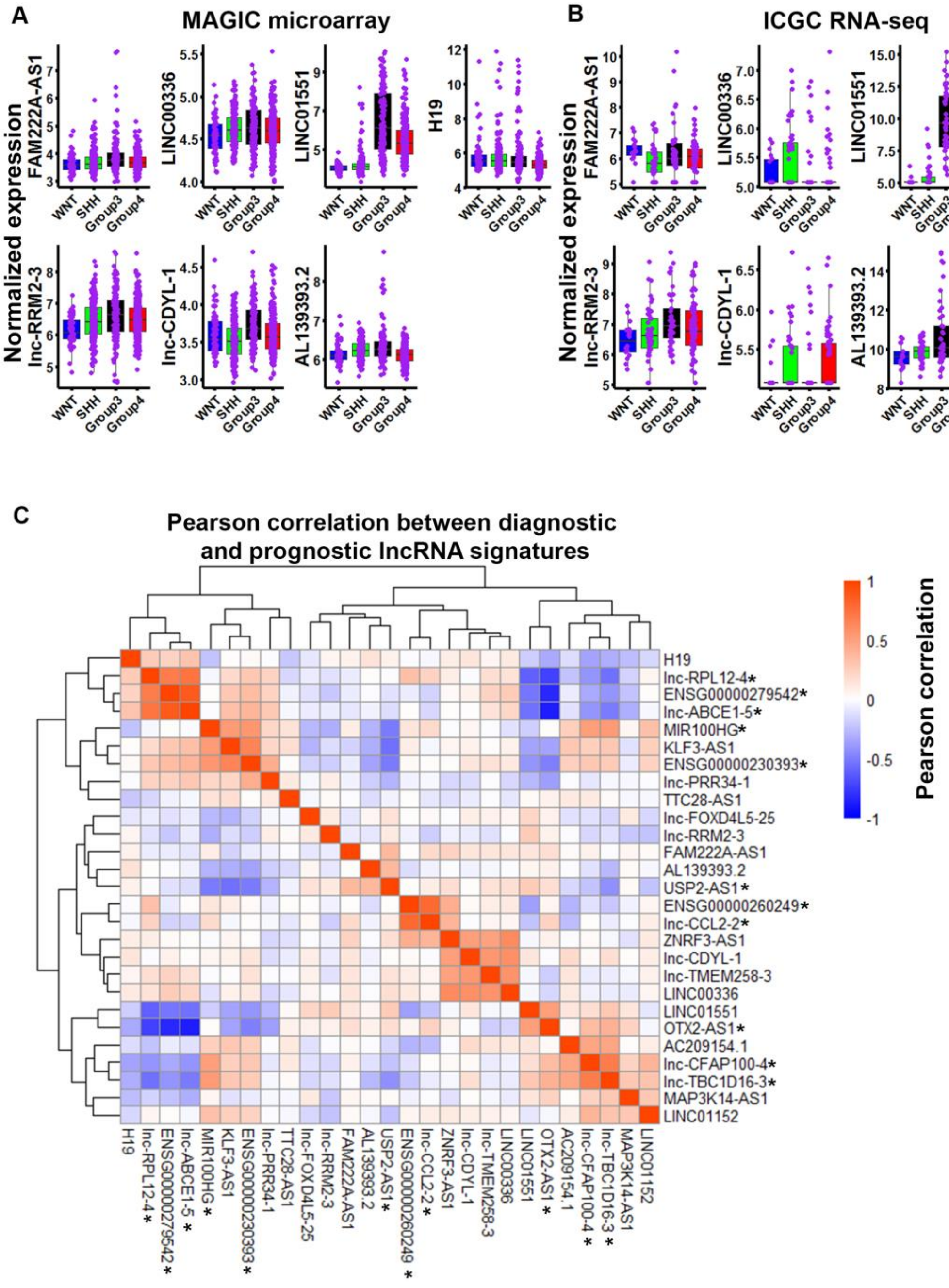
A
MAGIC microarray
B
ICGC RNA-seq
Normalized expression
FAM222A-AS1
LINC00336
LINC01551
H19
lnc-RRM2-3
lnc-CDYL-1
AL139393.2
WNT
SHH
Group3
Group4
C
Pearson correlation between diagnostic and prognostic lncRNA signatures
Pearson correlation
H19
lnc-RPL12-4*
ENSG00000279542*
lnc-ABCE1-5*
MIR100HG*
KLF3-AS1
ENSG00000230393*
lnc-PRR34-1
TTC28-AS1
lnc-FOXD4L5-25
lnc-RRM2-3
FAM222A-AS1
AL139393.2
USP2-AS1*
ENSG00000260249*
lnc-CCL2-2*
ZNRF3-AS1
lnc-CDYL-1
lnc-TMEM258-3
LINC00336
LINC01551
OTX2-AS1*
AC209154.1
lnc-CFAP100-4*
lnc-TBC1D16-3*
MAP3K14-AS1
LINC01152
